# Anaerobic derivates of mitochondria and peroxisomes in the free-living amoeba *Pelomyxa schiedti* revealed by single-cell genomics

**DOI:** 10.1101/2021.05.20.444135

**Authors:** Kristína Záhonová, Sebastian Cristian Treitli, Tien Le, Ingrid Škodová-Sveráková, Pavla Hanousková, Ivan Čepička, Jan Tachezy, Vladimír Hampl

**Author notes:** Corresponding authors (KZ); (VH).

## Abstract

*Pelomyxa schiedti* is a free-living amoeba belonging to the group Archamoebae, which encompasses anaerobes bearing mitochondrion-related organelles (MROs) – hydrogenosomes in free-living *Mastigamoeba balamuthi* and mitosomes in the human pathogen *Entamoeba histolytica*. Anaerobic peroxisomes, another adaptation to anaerobic lifestyle, were identified only recently in *M. balamuthi*. We found evidence for both these organelles in the single-cell-derived genome and transcriptome of *P. schiedti*, and corresponding vesicles were tentatively revealed in electron micrographs. *In silico* reconstructed MRO metabolism seems similar to that of *M. balamuthi* harboring respiratory complex II, electron-transferring flavoprotein, partial TCA cycle running presumably in reductive direction, pyruvate:ferredoxin oxidoreductase, [FeFe]-hydrogenases, glycine cleavage system, and sulfate activation pathway. The cell disposes with an expanded set of NIF enzymes for iron sulfur cluster assembly, but their localization remains unclear. Quite contrary, out of 67 predicted peroxisomal enzymes, only four were reported also in *M. balamuthi*, namely peroxisomal processing peptidase, nudix hydrolase, inositol 2-dehydrogenase, and D-lactate dehydrogenase. Other putative functions of peroxisomes could be pyridoxal 5I⍰-phosphate biosynthesis, amino acid and carbohydrate metabolism, and hydrolase activities. Future experimental evidence is necessary to define functions of this surprisingly enzyme-rich anaerobic peroxisome.

**Author summary:** A major part of the microbial diversity cannot be cultured in isolation, and so it escapes from traditional ways of investigation. In this paper, we demonstrate the successful approach for generating good-quality genome and transcriptome drafts from a peculiar amoeba *Pelomyxa schiedti* using single-cell methods. *P. schiedti* is a member of Archamoebae clade harboring microaerobic protists including a free-living *Mastigamoeba balamuthi* and a human parasite *Entamoeba histolytica*. Mitochondria and peroxisomes represent two organelles that are most affected during adaptation to microoxic or anoxic environments. Mitochondria are known to transform to anaerobic mitochondria, hydrogenosomes, mitosomes, and various transition stages in between, all of which encompass different enzymatic capacity. Anaerobic peroxisomes have been first noticed in *M. balamuthi*, but their function remained unclear for now. Data obtained in this study were used for revealing the presence and for the detailed functional annotations of anaerobic derivates of mitochondria and peroxisomes in *P. schiedti*, which were corroborated by transmission electron microscopy.

## Introduction

Transition to life in low oxygen environments requires significant modifications of cell biochemistry and organellar make up. Several lineages of protists have undergone such transitions and exemplify partly convergent solutions [1–3]. Mitochondria and peroxisomes have been most significantly remodeled in this process, as they are the key places of oxygen-dependent metabolism and oxygen detoxification.

Mitochondria are double-membrane-bound organelles, which have arisen from engulfment of a prokaryotic lineage related to alphaproteobacteria [2,4–6]. Since then, they have diverged into a range of categories [1] and plethora of transitional forms [7,8], collectively designated as mitochondrion related organelles (MROs), while only a single case of complete loss has been reported [9]. Substantial collection of typical mitochondrial functionalities, such as oxidative phosphorylation, carbon, amino acid and fatty-acid metabolism, iron-sulfur (FeS) cluster assembly, homeostasis, and apoptosis, has been reduced to various extent in MROs [10–12].

Peroxisomes are bound by a single membrane and characterized by a highly conserved set of proteins (peroxins) essential for their biogenesis [13,14]. The matrix content and consequently the repertoire of metabolic pathways is very variable reflecting high versatility of peroxisomal functions [15]. Most frequently, they possess oxidases reducing molecular oxygen to hydrogen peroxide (H_2_O_2_), and catalase for its detoxification. Not surprisingly, they are absent from most anaerobes, such as *Giardia* and *Trichomona*s [16]; however, anaerobic peroxisomes were recently reported from *Mastigamoeba balamuthi* [17].

Archamoebae represents a clade of microaerophilic protists nested within a broader group of predominantly aerobic amoebozoans [18,19] represented e.g. by *Dictyostelium discoideum*, known to bear a classical aerobic mitochondrion [20], or by their more distant amoebozoan relative *Acanthamoeba castellanii* (Centramoebida) with mitochondria potentially adapted to periods of anaerobiosis and exhibiting a highly complex proteome [12,21]. Small to almost inconspicuous MROs have been characterized in two Archamoebae, the parasitic *Entamoeba histolytica* and the free-living *M. balamuthi*. The only known function of *E. histolytica* mitosome is production and export of activated sulfate – phosphoadenosine-5’-phosphosulfate (PAPS) [22]. Metabolic capacity of *M. balamuthi* hydrogenosome is substantially broader involving pyruvate and amino acid metabolism, ATP production, and FeS cluster assembly [23–25]. Another adaptation of *M. balamuthi* to the low oxygen environment is represented by anaerobic peroxisomes that lack catalase and enzymes of β-oxidation of fatty acids but harbor several enzymes of pyrimidine and CoA biosynthesis and acyl-CoA and carbohydrate metabolism [17].

*Pelomyxa* is a free-living archamoeba distantly related to *M. balamuthi* [18], and so it represents valuable point for tracing the evolution of anaerobic adaptations. There is a single report on MROs in the giant species *P. palustris* [26] but their metabolism in unknown. Using methods of single-cell -omics and electron microscopy, we bring clear evidence for the presence of both MROs and peroxisomes in its smaller cousin *P. schiedti* [27].

## Results and discussion

### General features of assemblies

*P. schiedti* single-cell genome assembly of 52.4 Mb contained 5,338 scaffolds with an N50=51,552 bp (S1 Table) and 19,965 predicted proteins. We identified a single small subunit ribosomal RNA gene (18S rDNA). In the 18S phylogeny, *P. schiedti* was sister to other *Pelomyxa* species inside the Pelomyxidae clade (88% standard bootstrap) within a robust clade (94% standard bootstrap) of Archamoebae (Fig 1, S1 Fig). The decontaminated transcriptome assembly of 76.6 Mb comprised 43,993 contigs. BUSCO was used to estimate completeness of assemblies and to compare them to the *M. balamuthi* genome (S2 Fig, S1 Table). Transcriptome contained 83.2% of complete and 2.0% of fragmented BUSCO genes, while in the genome-derived proteome the proportions were 81.9% and 3.6%, respectively. With 82.8% complete and 3.0% fragmented genes [28], the completeness of *M. balamuthi* data was comparable. 36.0% of BUSCOs were duplicated in the transcriptome assembly, while only 8.6% in genomic, reflecting a higher number of contigs or presence of isoforms in the former. It should be noted that for non-model eukaryotes, which *Pelomyxa* certainly is, the BUSCO completeness is not expected to reach 100%, because some of the orthologues might be absent and/or diverged beyond recognition. Altogether, our analyses showed considerably high completeness of both assemblies.

**Fig 1.**
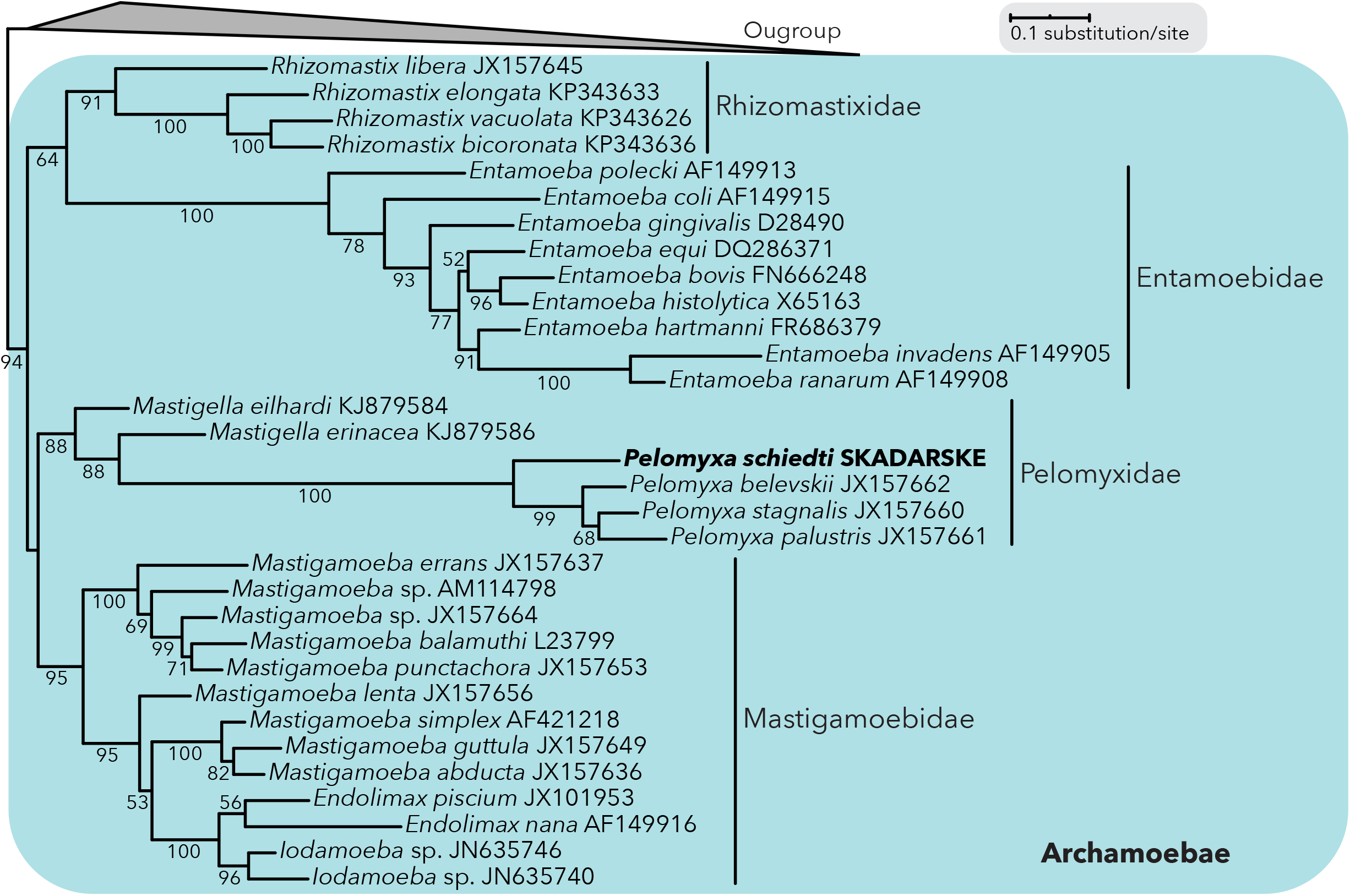
Phylogenetic analysis of amoebozoan 18S rDNA. The Maximum Likelihood tree places *Pelomyxa schiedti* in monophyletic Pelomyxidae group inside monophyletic Archamoebae. Standard bootstrap support values are shown when ≥ 50%. Outgroup was collapsed for simplicity (for full tree see S1 Fig).

*P. schiedti* genes encompass 149,016 introns (S1 Table), which accounts for an intron density of 7.46, almost twice higher than in *M. balamuthi* (3.74). While protists’ genomes have usually lower intron densities, several organisms in IntronDB [29] exhibit similar intron density as *Pelomyxa*, e.g., the choanoflagellates *Monosiga brevicolis* (6.53) and *Salpingoeca rosetta* (7.44), the chromerid *Vitrella brassicaformis* (7.45), or the chlorarachniophyte *Bigelowiella natans* (7.85). The vast majority of introns (98.41%) contained canonical GT-AG boundaries, 1.59% possessed GC-AG boundaries, and one an unusual GT-GG intron boundary (S1 Table). Similar frequencies of intron boundaries were observed in *M. balamuthi* (S1 Table and [28]).

### Putative MRO proteome

The major focus of this study was to reveal the presence and to characterize the putative proteomes of MRO and peroxisome of *P. schiedti*. We used a combined approach to search for proteins possibly involved in the MRO metabolism and biogenesis by: (i) retrieving homologues of MRO- or mitochondrion-targeted proteins of *E. histolytica, M. balamuthi*, and *A. castelanii*, and (ii) predicting N-terminal mitochondrial targeting sequence (NTS) by four tools. The resulting *in silico* predicted MRO proteome consists of 51 proteins (Fig 2, S2 Table) and provides functionalities described below.

**Fig 2.**
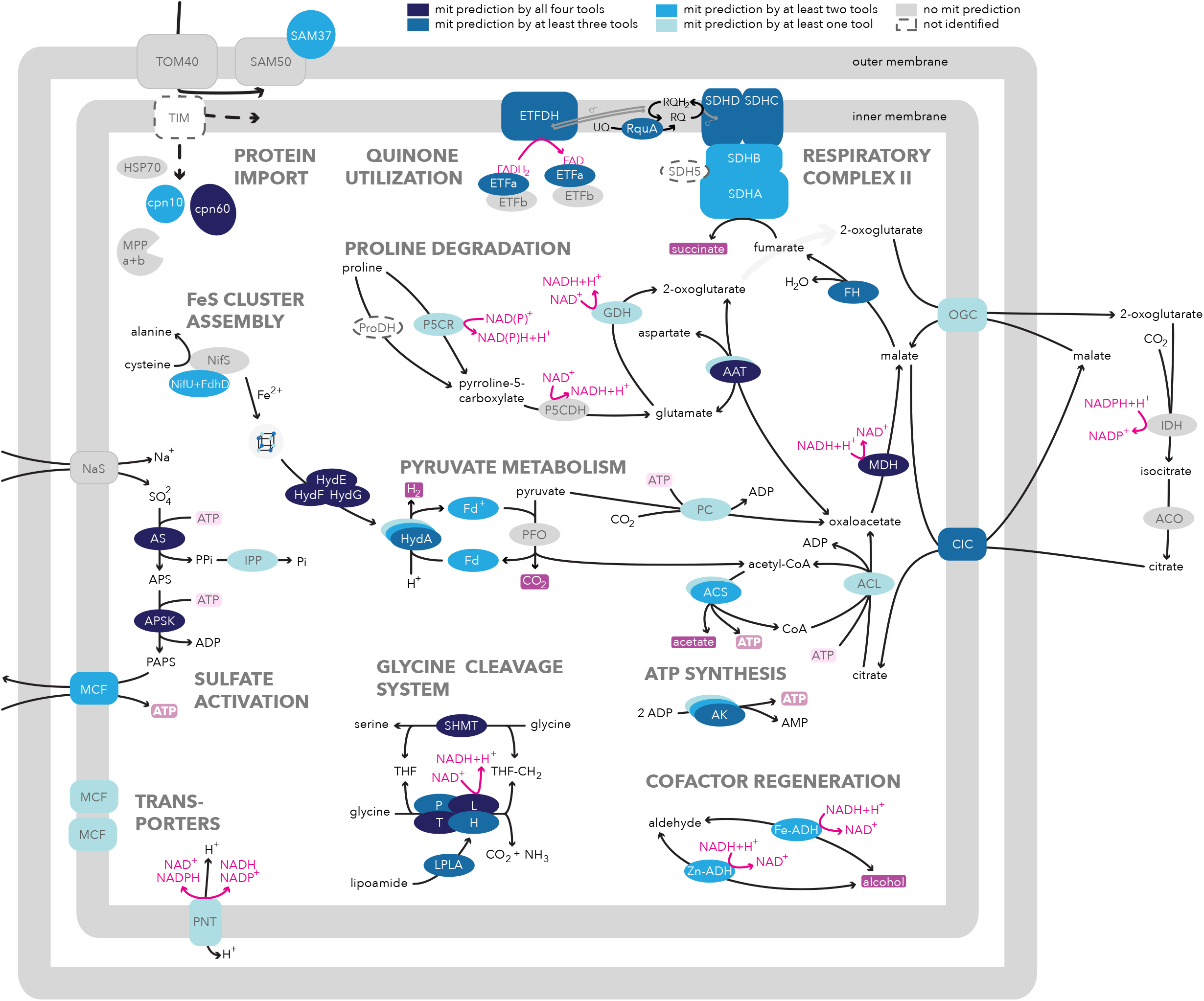
Overview of the Pelomyxa schiedti MRO metabolism. Proteins were identified by BLAST or HMMER searches and their intracellular localization was predicted by TargetP, PSORT II, MultiLoc2, and NommPred tools. Confidence of MRO localization is enhanced by shades of blue as explained in graphical legend above the scheme. Multiple copies of a protein are shown as overlapping ovals. Potential end-products are boxed in dark-fuchsia color. ATP production and consumption are highlighted by dark- and light-pink boxes around ATP, respectively. Abbreviations: AAT, aspartate alanine transferase; ACL, ATP-citrate lyase; ACO, aconitase; ACS, acetyl-CoA synthetase; AK, adenylate kinase; APS, adenosine-5’-phosphosulfate; APSK, adenosine-5’-phosphosulfate kinase; AS, ATP sulfurylase; cpn10, chaperonin 10; cpn60, chaperonin 60; CIC, citrate carrier; CoA, coenzyme A; ETFa, electron transferring flavoprotein subunit alpha; ETFb, electron transferring flavoprotein subunit beta; ETFDH, electron transferring flavoprotein dehydrogenase; Fe-ADH, iron-containing alcohol dehydrogenase; Fd, ferredoxin; FH, fumarase; GDH, glutamate dehydrogenase; H, GCSH protein; HSP70, heat shock protein 70; HydA, [FeFe]-hydrogenase; HydE, hydrogenase maturase; HydF, hydrogenase maturase; HydG, hydrogenase maturase; IDH, isocitrate dehydrogenase; IPP, inorganic pyrophosphatase; L, GCSL protein; D-LDH, D-lactate dehydrogenase; LPLA, lipoamide protein ligase; MCF, mitochondrial carrier family; MDH, malate dehydrogenase; MPP a+b, mitochondrial processing peptidase subunit alpha and beta; NaS, sodium/sulfate symporter; NifS, cysteine desulfurase; NifU+FdhD, scaffold protein + formate dehydrogenase accessory sulfurtransferase; OGC, 2-oxoglutarate carrier; P, GCSP protein; P5CDH, pyrroline-5-carboxylate dehydrogenase; P5CR, pyrroline-5-carboxylate reductase; PAPS, 3’-phosphoadenosine 5’-phosphosulfate; PC, pyruvate carboxylase; PFO, pyruvate:ferredoxin oxidoreductase; PNT, pyridine nucleotide transhydrogenase; ProDH, proline dehydrogenase; RQ, rodoquinone; RQH2, rhodoquinol; RquA, RQ methyltransferase; SAM, sorting and assembly machinery; SDH5, succinate dehydrogenase assembly factor; SDHA, succinate dehydrogenase subunit A; SDHB, succinate dehydrogenase subunit B; SDHC, succinate dehydrogenase subunit C; SDHD, succinate dehydrogenase subunit D; SHMT, serine hydroxymethyltransferase; T, GCST protein; THF, tetrahydrofolate; THF-CH2, N^5^,N^10^-methylenetetrahydrofolate; TOM/TIM, translocase of the outer/inner membrane; UQ, ubiquinone; Zn-ADH, zinc-containing alcohol dehydrogenase.

### Protein import machinery

Despite sensitive HMMER searching we identified only three subunits of the outer membrane translocase (TOM) and the sorting and assembly machinery (SAM) complexes — Tom40, Sam50, and Sam37 (Fig 2). All three proteins had corresponding domains predicted by InterProScan. Many homologues of the canonical opisthokont subunits are missing (S2 Table), as are all parts of the translocase of the inner membrane (TIM), and so the mechanism of protein import across this membrane remains unclear. The situation resembles other Archamoebae [23,28,30], suggesting that their translocons are either highly streamlined and/or contain highly divergent or lineage-specific subunits as reported from trichomonads or trypanosomes [31,32].

Enzymes involved in processing (matrix processing peptidase) and folding (chaperonins cpn10 and cpn60) are present. HSP70 was detected in 14 copies, none of them confidently predicted to mitochondrion (S2 Table). Phylogenetic analysis revealed a single MRO candidate (Pelo10550) branching sister to *M. balamuthi* mtHSP70 within the mitochondrial clade (S3 Fig). The other HSP70 paralogues fell into the ER or cytosolic clades, the latter being diversified in ten copies all forming robust clades with *M. balamuthi* sequences.

Although we have probably revealed only a fragment of the inventory needed for the protein import into the *P. schiedtii* MRO, the presence of the hallmarks—Tom40, Sam50, mtHSP70, and cpn60—conclusively shows that the MRO is truly present.

### Tricarboxylic acid cycle and electron transport chain

*P. schiedti* encodes four enzymes of the tricarboxylic acid (TCA) cycle possessing NTS (S2 Table) and catalyzing consecutive reactions. ATP citrate lyase (ACL) is typical for the reductive direction of TCA, while others, fumarate hydratase (fumarase/FH), malate dehydrogenase (MDH), and four subunits of the succinate dehydrogenase complex (SDH/complex II/CII), are common for both, oxidative and reductive TCA. The absence of CII subunit SDH5/SDHAF involved in the flavination of SDHA subunit [33] is likely common for Archamoebae as it is absent also from *M. balamuthi* [24].

Homologues of *A. castellanii* respiratory complexes were not identified, except for the aforementioned CII and a quinone-dependent electron-transferring flavoprotein (ETF; S2 Table). Both soluble subunits, alpha (ETFa) and beta (ETFb), and the membrane-bound ETF dehydrogenase (ETFDH), are present but only ETFDH and ETFa contain a recognizable NTS. It has been proposed in *M. balamuthi* that electrons may be transferred in an unknown direction between ETF and rhodoquinone (RQ), a quinone molecule with a lower electron potential than ubiquinone [3,34]. RQ is in *M. balamuthi* synthetized by a hydrogenosomal methyltransferase dubbed RquA [34], which was detected also in *P. schiedti* (S2 Table). RQ presence allows delivery of electrons to CII that could function as fumarate reductase [35] producing succinate, the putative end product of the partial reverted TCA in both Archamoebae [24], which may be secreted as in *Trypanosoma* [36].

### Pyruvate and ATP metabolism

Pyruvate is in aerobic mitochondria oxidatively decarboxylated to acetyl-coenzyme A (CoA) by the pyruvate dehydrogenase (PDH) complex. In most MROs, PDH is substituted by pyruvate:ferredoxin oxidoreductase (PFO), pyruvate:NADP^+^ oxidoreductase (PNO), or pyruvate formate lyase (PFL) [2]. We identified six copies of PFO and one copy of PNO in the *P. schiedti* genome, all without NTS (S2 Table). However, one of the *P. schiedti* PFOs was sister to one of the *M. balamuthi* putatively hydrogenosomal PFOs [24] (S4 Fig). We assume that this PFO homologue operates in *P. schiedti* MRO. Another pyruvate-metabolizing enzyme predicted to MRO is pyruvate carboxylase (PC; S2 Table) producing oxaloacetate [37], a substrate of MDH. In *M. balamuthi*, pyruvate may be produced by the activity of NAD^+^-dependent D-lactate dehydrogenase (D-LDH), of which one is present in hydrogenosome and the other in peroxisome [17,24]. *P. schiedti* bears only one homologue of D-LDH that is predicted to peroxisomes (S3 Table), thus pyruvate is likely imported to MRO from cytosol.

Two ATP-synthesizing enzymes are putatively present. Acetyl-CoA synthetase (ACS), enzyme converting acetyl-CoA to acetate, CoA, and ATP, was found in eight copies, four of which possessed a putative NTS. ATP may be formed also by the adenylate kinase (AK) catalyzing interconversion of adenine nucleotides. Three of the six AKs are putatively localized in the MRO (S2 Table). In this respect, the situation resembles *M. balamuthi* hydrogenosome [24]. Third putative source of ATP is the antiport against PAPS.

### Amino acid metabolism

Glycine cleavage system (GCS) is at least partially retained in many MROs [38]. It consists of four enzymes (H-, L-, T-, and P-protein) and methylates tetrahydrofolate (THF) while decomposing glycine into CO_2_ and ammonia. THF methylation is also provided by the serine hydroxymethyltransferase (SHMT) [39]. We identified all GCS enzymes and SHMT in *P. schiedti*, all with predicted NTS (Fig 2, S2 Table). L-protein was present in two copies with only one bearing NTS, similarly to *M. balamuthi*. The function of the second copy is unknown [24]. Lipoamide protein ligase (LPLA) necessary for lipoamide attachment to GCSH was present with NTS. The resulting N^5^,N^10^-methylenetetrahydrofolate (CH_2_ -THF) is an intermediate in one-carbon metabolism and cofactor for the synthesis of pyrimidines and methionine in both mitochondria and cytosol. Two cytosolic enzymes requiring this cofactor, B12-dependent methionine synthase and THF dehydrogenase/cyclohydrolase, were detected (S2 Table). Glycine can be produced in mitochondria from threonine by threonine dehydrogenase (TDH) and alpha-amino-beta-ketobutyrate CoA ligase (AKL) [40] but both proteins lack a recognizable NTS in *P. schiedti* (S2 Table). Consistently, TDH activity was measured only in the cytosolic fraction of *M. balamuthi* [24]. It is highly probable that this pathway operates in the cytosol of *P. schiedti* and glycine is imported to MRO.

To our surprise, we identified remnants of the proline degradation pathway presumably residing in *P. schiedti* MRO (Fig 2). In mitochondria, proline is usually degraded to glutamate by the function of proline dehydrogenase (ProDH) and pyrroline-5-carboxylate dehydrogenase (P5CDH) [41]. While ProDH is missing in *P. schiedti*, an alternative enzyme pyrroline-5-carboxylate reductase (P5CR) was predicted to be mitochondrion-targeted by one predictor (S2 Table). P5CDH is present in *P. schiedti* but lacks predictable NTS. Glutamate can be further metabolized to 2-oxoglutarate by glutamate dehydrogenase (GDH) [41], which is present and predicted to be mitochondrion-targeted also by one tool (S2 Table).

### Cofactor regeneration

NADH produced by GCS or during putative proline degradation would be in most mitochondria reoxidized by NADH dehydrogenases in the electron transport chain [41]. Since this is absent in *P. schiedti*, we explored other ways for regeneration of this cofactor. One possibility is fermentation of aldehydes to alcohols by alcohol dehydrogenases [42] putatively targeted to the MRO (S2 Table). Another option is the reductive partial TCA cycle running from citrate to succinate consuming not only NADH but also electrons from ETFDH *via* CII producing succinate [43]. Citrate or oxaloacetate are necessary to fuel this pathway. We identified a mitochondrial citrate carrier (CIC; S2 Table) which belongs to SLC25A family and is known to exchange malate for cytosolic citrate in cancer cells under low concentration of oxygen [44]. ACL produces acetyl-CoA and oxaloacetate from citrate on the expense of ATP [45]. Acetyl-CoA may become a substrate for anabolic reactions or be used by ACS to regenerate both ATP and CoA (Fig 2), while oxaloacetate may enter the reverse TCA cycle becoming substrate of MDH, regenerating NAD^+^. The malate pool is maintained also by 2-oxoglutarate carrier (OGC; S2 Table). In the cytosol, 2-oxoglutarate can be reductively carboxylated to replenish citrate [46]. Oxaloacetate may alternatively be produced from pyruvate by PC with ATP consumption or by the action of aspartate amino transferase (S2 Table). The latter enzyme may balance the ratio of 2-oxoglutarate + aspartate: oxalacetate + glutamate; however, the origin and fate of aspartate is unclear due to the absence of the glutamate-aspartate antiporter (S2 Table).

Pyridine nucleotide transhydrogenase (PNT) is predicted to MRO by a single predictor (S2 Table). PNT usually localizes in the inner mitochondrial membrane and pumps protons while transferring electrons between NADH and NADPH [47]. PNT is present in *M. balamuthi* and *E. histolytica* [23,48], however in *E. histolytica*, it was shown to localize outside mitosomes [49], which calls into question its MRO location in other Archamoebae.

MRO contains two additional electron sinks with unclear purpose. ETF and ETFDH proteins are known to use electrons from oxidation of fatty acids, which is absent in *P. schiedti* MRO. Finally, [FeFe]-hydrogenases uptake electrons from reduced ferredoxins and produce molecular hydrogen. Three of the six detected hydrogenases bear putative NTS. Hydrogenases contain catalytic H cluster and its maturation is dependent on maturases (HydE, HydF, and HydG) [50], which are all present and contain NTS (S2 Table). Reduced ferredoxin may originate from pyruvate oxidation.

### Iron-sulfur cluster assembly

Mitochondria usually house the iron-sulfur cluster assembly (ISC) pathway inherited from alphaproteobacteria serving for maturation of both, mitochondrial and cytosolic FeS proteins [51]. Some organisms, including Archamoebae, have replaced it by another machinery *via* horizontal gene transfer [25,52]. *M. balamuthi* bears two copies of the nitrogen fixation (NIF) system, both comprising NifS and NifU proteins. While one pair of NIFs operates in cytosol, the other localizes in the hydrogenosome [25]. In *E. histolytica*, only cytosolic copy has been retained [24].

In *P. schiedti* MRO, hydrogenases and their maturases HydE and HydF, SDH, ferredoxin, and PFO are putative clients for NIF system. We identified seven NifS and three NifU proteins, of which only NifU (Pelo10620) contained predicted NTS (S2 Table). Interestingly, this protein consists of a NifU N-terminal domain fused to a formate dehydrogenase accessory sulfurtransferase (FdhD) C-terminal domain (S5A Fig). The *Escherichia coli* FdhD transfers sulfur from IscS to formate dehydrogenase (FdhF) and is essential for its activity [53]. *P. schiedti* indeed encodes a FdhF homologue without NTS (S2 Table). In the NifU phylogeny (Fig 3A), the NifU domain of the fusion protein formed a long branch within a moderately supported (80% ultrafast bootstrap) clade of all Archamoebae NifUs. The other *P. schiedti* NifU sequences branched sister to hydrogenosomal and cytosolic *M. balamuthi* homologues. All three *P. schiedti* NifU sequences contained conserved cysteine residues (S5B Fig) necessary for their function [54].

**Fig 3.**
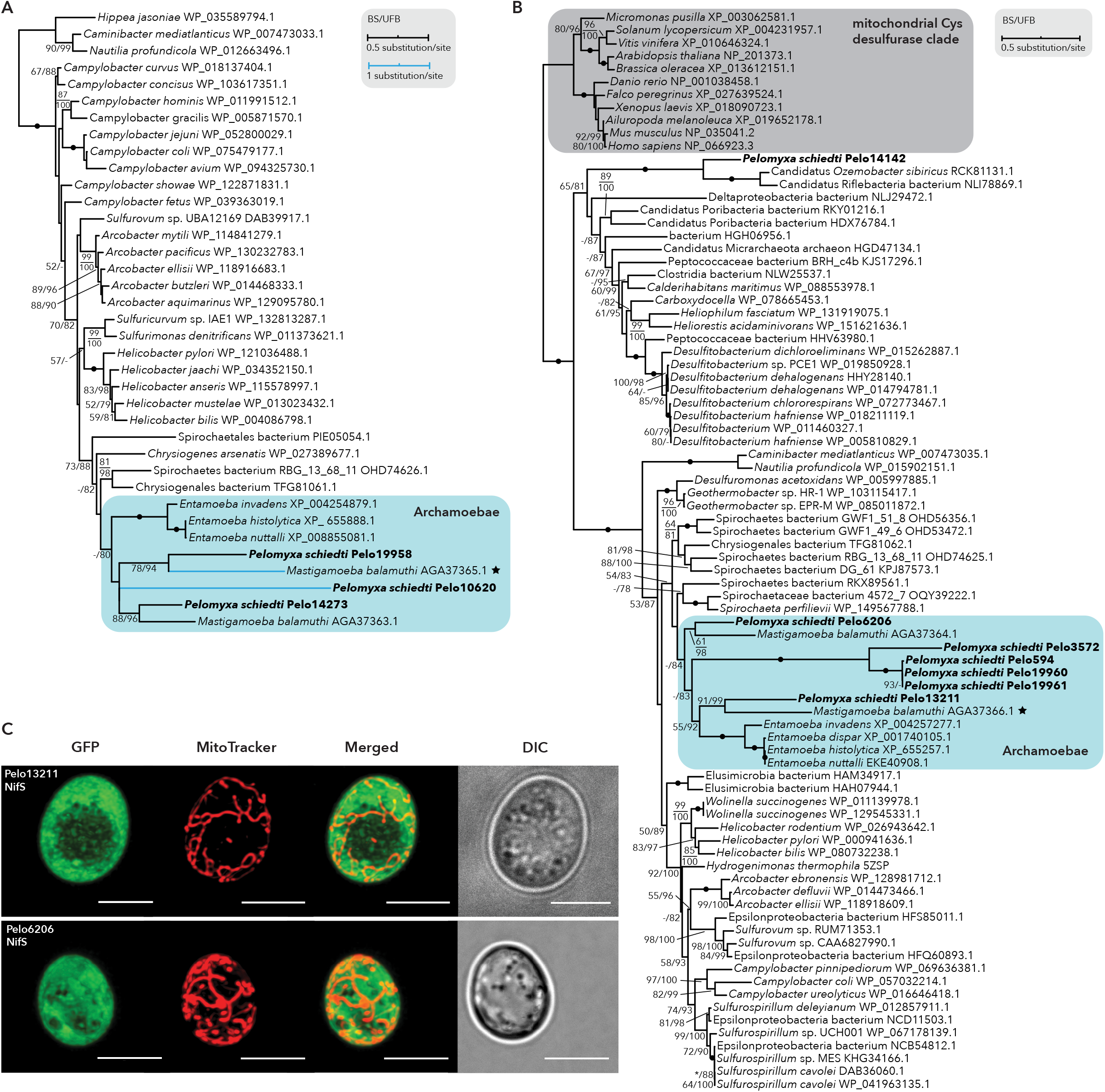
Analyses of NIF system components. (A-B) The Maximum Likelihood phylogenetic trees show that *Pelomyxa schiedti* possesses orthologues of hydrogenosomal and cytosolic NifU (A) and NifS (B) proteins from *Mastigamoeba balamuthi*. Hydrogenosomal proteins of *M. balamuthi* are marked with stars. The Maximum Likelihood tree was estimated with standard (BS) and ultrafast bootstrapping (UFB). The tree topology shown is from the ultrafast bootstrap analysis. Support values for <50% BS and <75% UFB are denoted by a dash (-), whereas an asterisk (*) marks a topology that does not exist in a particular analysis. Fully supported nodes are shown as black circles, while nodes that were not supported are without any value. (C) Heterologous expression of two NifS sequences of *Pelomyxa schiedti* showed their cytosolic localization. Proteins were expressed in *Saccharomyces cerevisiae* with a GFP-tag at their C-terminus. Mitochondria were stained with MitoTracker. DIC, differential interference contrast. Scale bar: 5 μm.

None of the seven NifS proteins was predicted to MRO (S2 Table). Two sequences were identical but incomplete at their C-termini and could not be completed by read mapping or PCR amplification. In the phylogenetic analysis (Fig 3B), Pelo13211 and Pelo6206 branched sister to the hydrogenosomal and cytosolic *M. balamuthi* sequences, respectively. Pelo14142 was sister to candidate Riflebacteria species and the last four formed a long branch nested within the Archamoebae clade. All amino acid residues required for function [55] were present in both *P. schiedti* sequences that were sister to *M. balamuthi* (S5C Fig).

The phylogenetic pattern offers an elegant hypothesis in which NifS Pelo6206 and NifU Pelo14273 act in the cytosol, while NifS Pelo13211 and NifU Pelo19958 in MRO. The remaining NifS copies might be functional partners of the NifU-FdhD fusion protein (Pelo10620). Surprisingly, our experiments with heterologous localization of the MRO and cytosolic NifS candidates in *Saccharomyces cerevisiae* revealed cytosolic localization of both (Fig 3C), leaving the question of the FeS cluster assembly in *P. schiedti* MRO unresolved.

### Sulfate activation pathway

Sulfate activation pathway produces PAPS necessary for sulfolipid synthesis [22]. It is present in *E. histolytica* [22,56] and *M. balamuthi* [24] MROs, and we identified all of its components also in *P. schiedti* (Fig 2, S2 Table). The pathway requires two transporters. A sodium/sulfate symporter (NaS) is necessary for substrate delivery, however, its homologues in *P. schiedti* (S2 Table) are unrelated to *E. histolytica* mitosomal NaS [22] (S6A Fig) yielding their role unclear. The PAPS exporter belongs to the mitochondrial carrier family (MCF) and, indeed, one of *P. schiedti* MCF proteins branched sister to a clade of PAPS transporters of *E. histolytica* and *M. balamuthi* [28,57] (S6B Fig). As this transporter exchanges PAPS to ATP, it plays role in supplementing the ATP pool in MRO, yet cannot provide a net ATP gain, because two ATP molecules are required for production of one PAPS.

### Anaerobic peroxisomes

We have also investigated the presence of anaerobic peroxisomes, which were recently characterized in *M. balamuthi* [17]. *P. schiedti* encodes genes for 13 proteins required for peroxisome biogenesis (peroxins, Pexs) strongly suggesting presence of peroxisomes. Identified peroxins include Pex5 and Pex7 required for the recognition of peroxisomal targeting signal 1 and 2 (PTS1 and PTS2), respectively, Pex13 and 14 mediating the protein import, Pex1, 2, 6, 10, and 12, which are receptor-recycling peroxins, Pex3, 16, and 19 involved in protein import to the peroxisomal membrane, and Pex11 participating in the peroxisome fission (S3 Table). Prediction of putative peroxisomal matrix proteins based on the PTS1/PTS2 presence revealed 67 candidates (S3 Table). Interestingly, only four candidates were previously found in anaerobic peroxisomes of *M. balamuthi* that include peroxisomal processing peptidase (PPP), inositol 2-dehydrogenase, nudix hydrolase, and D-lactate dehydrogenase, all with clear support for localization in *P. schiedti* peroxisomes (S3 Table). Unlike in *M. balamuthi, P. schiedti* peroxisomes possibly contain pyridoxamine 5’-phosphate oxidase (PNPO) that utilizes molecular oxygen as an electron acceptor to catalyze the last step of the pyridoxal 5IZ-phosphate (PLP) biosynthesis with concomitant formation of ammonia and H_2_O_2_. The presence of PNPO raises a question how H_2_O_2_ is detoxified as typical antioxidant enzymes, such as catalase and peroxidase, are not present. However, H_2_O_2_ could be decomposed also nonenzymatically by antioxidants, such as 2-oxoglutarate, in which the ketone group of the α-carbon atom reacts with H_2_O_2_ to form succinate, CO_2_, and water [58]. *P. schiedti* contains a putative glutamate dehydrogenase that may produce 2-oxoglutarate and that possesses –SKL triplet, a typical PTS1. However, the peroxisomal targeting was not supported by the PTS predictor, which considers twelve C-terminal amino acid residues. The other proteins with predicted peroxisomal localization include several enzymes of amino acid synthesis and degradation, carbohydrate metabolism and hydrolases, however without clear biochemical context. More experimental studies are required to verify predicted localizations and to delineate function of peroxisomes in *P. schiedti*.

### Electron microscopy

Finally, we were interested whether the two organelles described by the genomic data can be visualized by microscopy. Careful inspection of electron micrographs, indeed, revealed two populations of small vesicles, one presumably bounded by a double membrane while the other by a single (Fig 4). We ascribe them to putative MROs and peroxisomes *in silico* characterized in this work but leave the confirmation for further studies.

**Fig 4.**
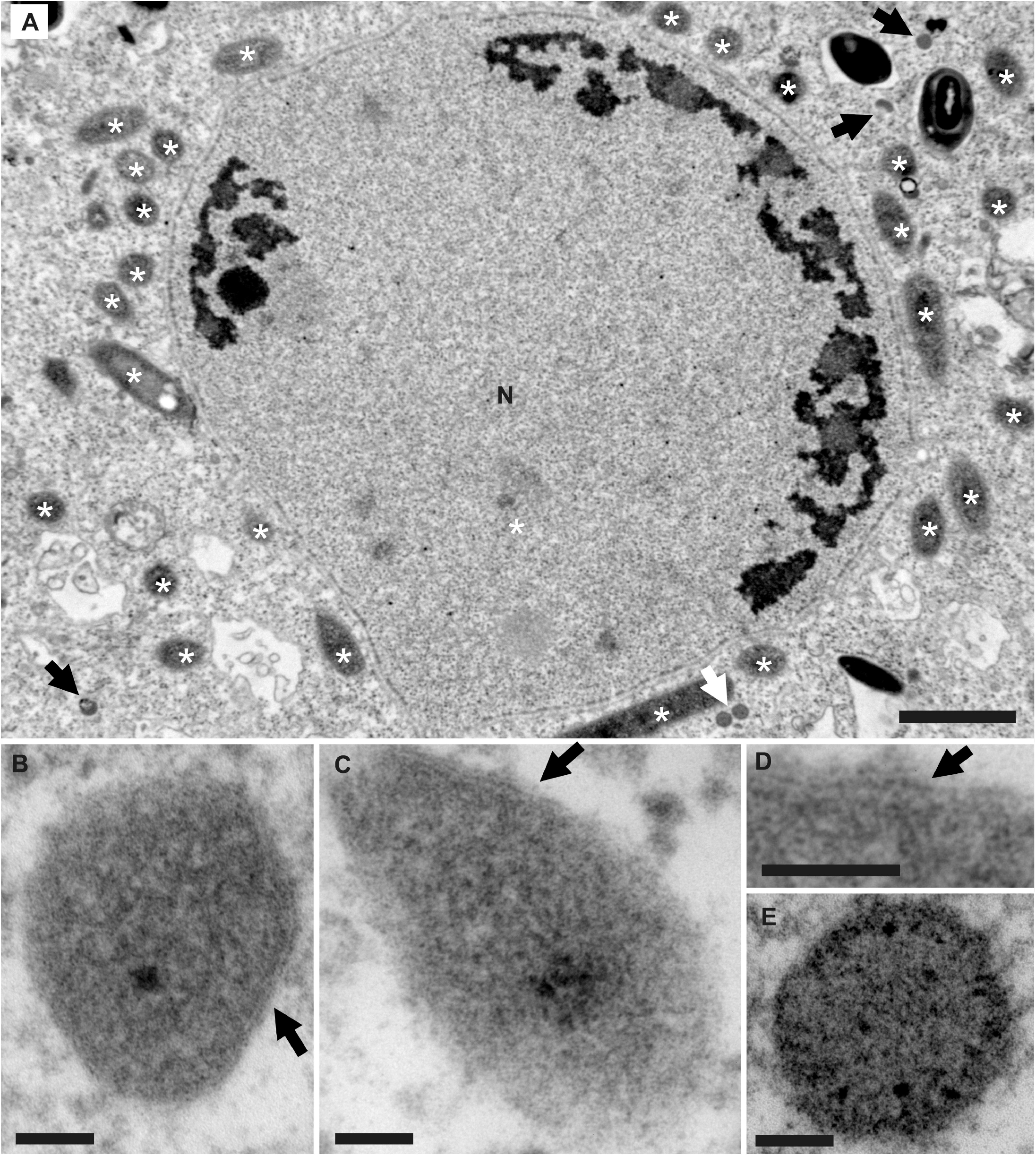
Transmission electron micrograph of Pelomyxa schiedti, ultra-thin sections. (A) The nuclear area. N, nucleus; black arrow, putative mitochondrion-related organelle; white arrow, small dense body (putative peroxisome); asterisk (*), prokaryotic endosymbiont. (B-C) High magnification of putative mitochondrion-related organelle; black arrow, bounding double membrane. (D) Detail of the bounding double membrane. (E) High magnification of the small dense body (putative peroxisome). Scale bars: 400 nm for (A); 50 nm for (B-E).

## Conclusions

Our bioinformatic survey of the putative proteome of *Pelomyxa schiedti* MRO revealed several interesting insights and opened many questions for further investigation of this amoeba. Most importantly, *P. schiedti* clearly does harbor an MRO with a very streamlined or lineage specific set of protein translocases, and peroxisomes with a set of 13 soluble and membrane associated peroxins. Our *in silico* predictions showed that the MRO provides the cell with the synthesis of PAPS, contains glycine cleavage system, [FeFe]-hydrogenase, and likely also a part of a TCA cycle running in reverse direction from citrate enabling concomitantly the production of acetyl-CoA. The electron transport chain is reduced to complex II and electron-transferring flavoprotein dehydrogenase, and possibly uses rhodoquinone as the electron transporter. We predict that the source of reduced ferredoxin for [FeFe]-hydrogenase comes from pyruvate. The situation with the FeS cluster assembly in this amoeba seems rather complex as it contains the most diverse set of NIF pathway proteins of all previously investigated Archamoebae. These proteins very likely provide parallel FeS synthesis in MRO and cytoplasm, but in addition to this, some may be involved in the activation of formate dehydrogenase as seen in some prokaryotes. *P. schiedti* anaerobic peroxisomes, similarly to *M. balamuthi*, lack enzymes of β-oxidation of fatty acids and catalase. Although the function of these peroxisomes needs to be clarified, the set of predicted enzymes suggested significant metabolic diversity between the two amoebae as well as from their aerobic counterparts.

## Materials and methods

### Cell culture

Polyxenic (and polyeukaryotic) culture of *Pelomyxa schiedti* strain SKADARSKE was maintained in Sonneborn’s *Paramecium* medium [59] as described previously [27].

### Genome and transcriptome sequencing and assembly

Genome sequencing was performed from whole genome amplified DNA (WGA). Individual cells were picked by micromanipulation and washed twice in Trager U media [60]. Genomic DNA was amplified using Illustra Single Cell GenomiPhi DNA Amplification Kit (GE Healthcare Life Sciences) according to the manufacturer’s protocol and purified using ethanol precipitation. Presence of the eukaryotic DNA was confirmed by amplification of a partial actin gene using specific primers (S4 Table). Sequencing libraries from seven positive samples were prepared using Illumina TruSeq DNA PCR-Free kit (Illumina). Samples Pelo2 and Pelo5 were sequenced on Illumina MiSeq (2×300 bp; Genomic Core facility, Faculty of Science) and Nanopore (Oxford Nanopore Technologies), samples P1 – P5 on Illumina HiSeq X (Macrogen Inc.). The Nanopore library was prepared using Oxford Nanopore Technologies ligation sequencing kit (SQK-LSK108) from 4 μg of T7 endonuclease I (New England Biolabs) treated WGA. Sequencing was performed using a R9.4.1 Spot-On Flow cell (FLO-MIN106) for 48 hours.

For transcriptome sequencing, single-cells of *P. schiedti* were washed twice in Trager U and amplification by 19 cycles was performed [61]. Five libraries were prepared using Nextera XT DNA Library preparation Kit (Illumina) and sequenced on Illumina MiSeq (PE 2×300bp; Genomic Core facility, Faculty of Science).

Raw Illumina DNA- and RNA-Seq reads were quality and adapter trimmed using BBDuk v36.92 (part of BBTools suite: https://jgi.doe.gov/data-and-tools/bbtools/). Firstly, individual single-cell genome assemblies for Pelo2, Pelo5, and P1 – P5 were generated with SPAdes v3.11.1 [62] using single-cell (--sc) mode and a k-mer size of 127. As the 18S rDNAs extracted from individual assemblies were identical, all reads (i.e., Illumina HiSeqs and MiSeq, and Nanopore) were assembled together by SPAdes v3.11.1 using --sc and k-mers of 21, 33, 55, 77, 99, 121. The resulting assembly was binned and decontaminated using tetraESOM [63] and a BLASTing strategy described previously [64]. The final assembly was scaffolded using P_RNA_scaffolder [65]. Prediction was done using Augustus v3.3.1 [66], and further improved by PASA and EVM [67] using the transcriptomic data. RNA-Seq reads were assembled using Trinity v2.6.5 [68] with default parameters, and contaminants were removed by BLASTing against the decontaminated genome assembly. RNA-Seq reads were mapped to the transcriptome using Bowtie2 v2.3.0 [69] and to the genome using HISAT2 v2.0.5 [70]. Genome and transcriptome completeness were assessed using BUSCO v3 with the eukaryota_odb9 dataset [71].

### Sequence searches and localization predictions

Proteins predicted to localize in *M. balamuthi* hydrogenosome, *E. histolytica* mitosome, and *A. castellanii* mitochondria served as queries in BLAST v2.6.0+ [72] searches through *P. schiedti* assemblies. Sensitive searches for components of TOM/TIM machinery were done using HMMER v3.3 [73]. Protein domains were predicted by InterProScan [74] implemented in Geneious Prime v2020.2.3 [75].

Potentially mitochondrion-targeted proteins were identified using TargetP v2 [76], PSORT II [77], MultiLoc2 [78], and NommPred [79] tools. Since *P. schiedti* does not harbor plastid, the plant setting from MultiLoc2 was omitted and NommPred was used in the MRO and in the *Dictyostelium* settings [19]. A protein was considered as mitochondrial if predicted by at least one setting of MultiLoc2 or NommPred.

Peroxins were identified by BLAST searches using *M. balamuthi* queries. Peroxisomal matrix proteins were predicted by searching for peroxisomal targeting signals (PTS). The tripeptides SRI and [SAP][KR][LM] (excluding AKM, PKM, and PRM) were used to search for the C-terminal PTS1. Proline at position -3 and methionine at position -1 were included based on experimental verification in *M. balamuthi* [17]. Two nanopeptides R[LI](x5)HL were used for N-terminal PTS2 searches [80]. All putative transmembrane proteins determined by TMHMM Server v2.0 [81] were filtered out. PTS1 candidates were submitted to the PTS1 Predictor using GENERAL function [82] evaluating twelve C-terminal residues.

### Phylogenetic analyses

An 18S rRNA gene dataset was aligned by MAFFT v7 [83] server with the G-INS-i algorithm at default settings and manually edited in BioEdit v7.0.4.1 [84] resolving 1,437 positions. Phylogenetic tree was constructed using Maximum-Likelihood in RAxML v8.0.0 [85] under the GTRGAMMAI model, 100 starting trees, and 1,000 bootstrap pseudoreplicates.

For selected proteins, datasets were aligned by MAFFT v7.313 [83], trimmed by trimAl v1.4 [86] and Maximum-Likelihood trees were inferred by IQ-TREE v1.6.8 [87] using the posterior mean site frequency method [88], LG+C20+F+G model, with the guide tree inferred under LG+F+G. Branch supports were obtained by the ultrafast bootstrap approximation [89].

### Immunofluorescence analysis

NifS genes (Pelo6206 and Pelo13211) were amplified from cDNA using specific primers (S4 Table) and PrimeSTAR^®^ Max DNA Polymerase (Takara Bio Inc.) premix, cloned into pUG35 vector containing C-terminal green fluorescence protein (GFP), and transformed to *S. cerevisiae* strain YPH499 using the lithium acetate method [90]. Transformants were grown on selective medium without uracil (SD-URA) at 30 °C. For localization, transformed cells were incubated with MitoTracker Red CMXRos (1:10,000; Thermo Fisher Scientific) for 10 minutes, followed by two washes with PBS, and mounted in 1% low-melting agarose and imaged using a Leica SP8 confocal microscope. Deconvolution was performed using Huygens Professional v17.10 and ImageJ v1.50b.

### Transmission electron microscopy

A grown culture of *P. schiedti* was pelleted by centrifugation and fixed one hour on ice with 2.5% glutaraldehyde in 0.1 M cacodylate buffer (pH 7.2). After washing in 0.1 M cacodylate buffer, the cells were postfixed one hour on ice with 1% OsO_4_. After washing with distilled water, the fixed cells were dehydrated in a graded series of ethanol, transferred to acetone, and embedded in EPON resin. Ultrathin sections were prepared on an ultramicrotome (Reichert-Jung Ultracut E) with a diamond knife. Sections were stained with uranyl acetate and lead citrate and examined using JEOL 1011 transmission electron microscope.

## Data availability

The raw sequencing data are available at NCBI (https://www.ncbi.nlm.nih.gov/) as BioProject PRJNA672820. Final assemblies are available from Zenodo at https://zenodo.org/record/4733726#.YI_RPWYza3I.

## Author contributions

SCT cultured cells, prepared sequencing libraries, and performed Nanopore sequencing. KZ and SCT assembled genome and transcriptome, and predicted proteins. IČ provided the culture of *P. schiedti* and conducted 18S rDNA phylogeny. KZ performed most of the bioinformatic and phylogenetic analyses. KZ, IŠ-S, JT, and VH analyzed the metabolism of the mitochondrion-related organelle. TL and JT analyzed the peroxisomal metabolism. PH performed the microscopic observations. JT and VH supervised the project. KZ, JT, and VH wrote the manuscript. All authors contributed to the editing of the final manuscript.

## Acknowledgements

We thank Dr. Zoltán Füssy (Charles University, BIOCEV, Prague, Czech Republic) and Dr. Courtney W. Stairs (Lund University, Lund, Sweden) for helpful discussions and advice.

## Supporting information

**S1 Fig. Phylogenetic analysis of amoebozoan 18S rDNA**. The Maximum Likelihood tree places *Pelomyxa schiedti* in monophyletic Pelomyxidae group inside monophyletic Archamoebae. Standard bootstrap support values are shown when ≥ 50%.

**S2 Fig. BUSCO analysis of the *Pelomyxa schiedti* transcriptome and predicted proteins**. Completeness of *P. schiedti* datasets were assessed using the odb9_eukaryota dataset and compared with completeness of predicted proteins from *Mastigamoeba balamuthi*.

**S3 Fig. Phylogenetic analysis of HSP70 proteins**. The Maximum Likelihood phylogenetic tree documents that one of the *Pelomyxa schiedti* HSP70 sequence is related to mitochondrial orthologues from other eukaryotes. Ultrafast bootstrap support values are shown when ≥ 75%.

**S4 Fig. Phylogenetic analysis of PFO enzymes**. The Maximum Likelihood phylogenetic tree identified a PFO version putatively operating in *Pelomyxa schiedti* MRO. Hydrogenosomal PFO copies of *Mastigamoeba balamuthi* are marked with stars. Number in parenthesis shows number of species in the collapsed clade. Ultrafast bootstrap support values are shown when ≥ 75%.

**S5 Fig. Sequences of *Pelomyxa schiedti* components of NIF system**. (A) The diagram depicts *P. schiedti* protein Pelo10620 composed of a NifU N-terminal domain fused to a FdhD (formate dehydrogenase accessory sulfurtransferase) C-terminal domain as determined by InterProScan. (B-C) Sequence alignment of NifU (B) and NifS (C) proteins from *P. schiedti* and *Mastigamoeba balamuthi* in comparison with bacterial homologues from *Thermotoga maritima*. The amino acid residues necessary for the function of NifU and NifS are labeled according to the legend.

**S6 Fig. Phylogenetic analysis of transporters involved in the sulfate activation pathway**. (A) The phylogenetic analysis did not resolve which one of the sodium/sulfate symporters of *Pelomyxa schiedti* is related to the *Entamoeba histolytica* mitosomal transporter. (B) The Maximum Likelihood phylogenetic tree confirms one *P. schiedti* transporter as PAPS (3’-phosphoadenosine 5’-phosphosulfate) transporter, while two others belong to a broader mitochondrial carrier family of transporters. Experimentally proven mitosomal transporters of *E. histolytica* are marked with stars. Ultrafast bootstrap support values are shown when ≥ 75%.

**S1 Table: Statistics of *Pelomyxa schiedti* assemblies** were compared with those of *Mastigamoeba balamuthi*.

**S2 Table: Proteins targeted to MRO of *Pelomyxa schiedti***. Localization of proteins was predicted by several tools, as listed in columns E - J. Mitochondrial predictions are highlighted by white font on blue background. Column K shows final inferred prediction of localization. mit, mitochondrial; cyt, cytosolic; SP, signal peptide; ER, endoplasmic reticulum; nuc, nuclear; extracell, extracellular; sec, secretory system; perox, peroxisomal; PM, plasma membrane; Other, other localization; -, protein not localized in MRO; +, protein localized in MRO; +?, protein localized in MRO with low confidence.

**S3 Table. Proteins required for peroxisome biogenesis and targeted to peroxisome**. Proteins identified in *Mastigamoeba balamuthi* are highlighted by white font on blue background. Proteins were considered peroxisome-targeted, if they contained PTS1 (SRI or [SAP][KR][LM]) or PTS2 motif (R[LIV](x5)HL), and/or were predicted by PTS1 predictor [82].

**S4 Table: Primers used in this study**.

## References

1. Müller M, Mentel M, van Hellemond JJ, Henze K, Woehle C, Gould SB, et al. Biochemistry and evolution of anaerobic energy metabolism in eukaryotes. Microbiol Mol Biol Rev. 2012;76: 444–495.

2. Roger AJ, Muñoz-Gómez SA, Kamikawa R. The origin and diversification of mitochondria. Curr Biol. 2017;27: R1177–R1192.

3. Gawryluk RMR, Stairs CW. Diversity of electron transport chains in anaerobic protists. Biochim Biophys Acta - Bioenerg. 2021;1862: 148334.

4. Zaremba-Niedzwiedzka K, Caceres EF, Saw JH, Bäckström Di, Juzokaite L, Vancaester E, et al. Asgard archaea illuminate the origin of eukaryotic cellular complexity. Nature. 2017;541: 353–358.

5. Sagan L. On the origin of mitosing cells. J Theor Biol. 1967;14: 255–274.

6. Martijn J, Vosseberg J, Guy L, Offre P, Ettema TJG. Deep mitochondrial origin outside the sampled alphaproteobacteria. Nature. 2018;557: 101–105.

7. Gawryluk RMR, Kamikawa R, Stairs CW, Silberman JD, Brown MW, Roger AJ. The earliest stages of mitochondrial adaptation to low oxygen revealed in a novel rhizarian. Curr Biol. 2016;26: 2729–2738.

8. Leger MM, Kolisko M, Kamikawa R, Stairs CW, Kume K, Čepička I, et al. Organelles that illuminate the origins of Trichomonas hydrogenosomes and Giardia mitosomes. Nat Ecol Evol. 2017;1: 0092.

9. Karnkowska A, Vacek V, Zubáčová Z, Treitli SC, Petrželková R, Eme L, et al. A eukaryote without a mitochondrial organelle. Curr Biol. 2016;26: 1274–1284.

10. Panigrahi AK, Ogata Y, Zíková A, Anupama A, Dalley RA, Acestor N, et al. A comprehensive analysis of Trypanosoma brucei mitochondrial proteome. Proteomics. 2009;9: 434–450.

11. Lee CP, Taylor NL, Harvey Millar A. Recent advances in the composition and heterogeneity of the Arabidopsis mitochondrial proteome. Front Plant Sci. 2013;4: 4.

12. Gawryluk RMR, Chisholm KA, Pinto DM, Gray MW. Compositional complexity of the mitochondrial proteome of a unicellular eukaryote (Acanthamoeba castellanii, supergroup Amoebozoa) rivals that of animals, fungi, and plants. J Proteomics. 2014;109: 400–416.

13. Žárský V, Tachezy J. Evolutionary loss of peroxisomes - not limited to parasites. Biol Direct. 2015;10: 74.

14. Schlüter A, Fourcade S, Ripp R, Mandel JL, Poch O, Pujol A. The evolutionary origin of peroxisomes: An ER-peroxisome connection. Mol Biol Evol. 2006;23: 838–845.

15. Gabaldón T. Peroxisome diversity and evolution. Philos Trans R Soc B Biol Sci. 2010;365: 765–773.

16. Gabaldón T, Ginger ML, Michels PAM. Peroxisomes in parasitic protists. Mol Biochem Parasitol. 2016;209: 35–45.

17. Le T, Žárský V, Nývltová E, Rada P, Harant K, Vancová M, et al. Anaerobic peroxisomes in Mastigamoeba balamuthi. Proc Natl Acad Sci U S A. 2020;117: 2065–2075.

18. Pánek T, Zadrobílková E, Walker G, Brown MW, Gentekaki E, Hroudová M, et al. First multigene analysis of Archamoebae (Amoebozoa: Conosa) robustly reveals its phylogeny and shows that Entamoebidae represents a deep lineage of the group. Mol Phylogenet Evol. 2016;98: 41–51.

19. Kang S, Tice AK, Spiegel FW, Silberman JD, Pánek T, Cepicka I, et al. Between a pod and a hard test: The deep evolution of Amoebae. Mol Biol Evol. 2017;34: 2258–2270.

20. Pearce XG, Annesley SJ, Fisher PR. The Dictyostelium model for mitochondrial biology and disease. Int J Dev Biol. 2019;63: 497–508.

21. Leger MM, Gawryluk RMR, Gray MW, Roger AJ. Evidence for a hydrogenosomal-type anaerobic ATP generation pathway in Acanthamoeba castellanii. PLoS One. 2013;8: e69532.

22. Mi-ichi F, Yousuf MA, Nakada-Tsukui K, Nozaki T. Mitosomes in Entamoeba histolytica contain a sulfate activation pathway. Proc Natl Acad Sci. 2009;106: 21731–21736.

23. Gill EE, Diaz-Triviño S, Barberà MJ, Silberman JD, Stechmann A, Gaston D, et al. Novel mitochondrion-related organelles in the anaerobic amoeba Mastigamoeba balamuthi. Mol Microbiol. 2007;66: 1306–1320.

24. Nývltová E, Stairs CW, Hrdý I, Rídl J, Mach J, Pačes J, et al. Lateral gene transfer and gene duplication played a key role in the evolution of Mastigamoeba balamuthi hydrogenosomes. Mol Biol Evol. 2015/01/07. 2015;32: 1039–1055.

25. Nývltová E, Šuták R, Harant K, Šedinová M, Hrdy I, Paces J, et al. NIF-type iron-sulfur cluster assembly system is duplicated and distributed in the mitochondria and cytosol of Mastigamoeba balamuthi. Proc Natl Acad Sci U S A. 2013;110: 7371–7376.

26. Seravin L, Goodkov A. Cytoplasmic microbody-like granules of the amoeba Pelomyxa palustris. Tsitologiya. 1987;29: 600–603.

27. Zadrobílková E, Walker G, Čepička I. Morphological and molecular evidence support a close relationship between the free-living Archamoebae Mastigella and Pelomyxa. Protist. 2015;166: 14–41.

28. Žárský V, Klimeš V, Pačes J, Vlček Č, Hradilová M, Beneš V, et al. The Mastigamoeba balamuthi genome and the nature of the free-living ancestor of Entamoeba. Mol Biol Evol. 2021;msab020: .

29. Wang D, Hancock J. IntronDB: A database for eukaryotic intron features. Bioinformatics. 2019;35: 4400–4401.

30. Dolezal P, Dagley MJ, Kono M, Wolynec P, Likić VA, Foo JH, et al. The essentials of protein import in the degenerate mitochondrion of Entamoeba histolytica. PLoS Pathog. 2010;6: e1000812.

31. Schneider A. Evolution of mitochondrial protein import - Lessons from trypanosomes. Biol Chem. 2020;401: 663–676.

32. Makki A, Rada P, Žárský V, Kereïche S, Kováčik L, Novotný M, et al. Triplet-pore structure of a highly divergent TOM complex of hydrogenosomes in Trichomonas vaginalis. PLoS Biol. 2019;17: e3000098.

33. Hao H-X, Khalimonchuk O, Schraders M, Dephoure N, Bayley J-P, Kunst H, et al. SDH5, a gene required for flavination of succinate dehydrogenase, is mutated in paraganglioma. Science. 2009;325: 1139–1142.

34. Stairs CW, Eme L, Muñoz-Gómez SA, Cohen A, Dellaire G, Shepherd JN, et al. Microbial eukaryotes have adapted to hypoxia by horizontal acquisitions of a gene involved in rhodoquinone biosynthesis. Elife. 2018;7: e34292.

35. Castro-Guerrero NA, Jasso-Chávez R, Moreno-Sánchez R. Physiological role of rhodoquinone in Euglena gracilis mitochondria. Biochim Biophys Acta. 2005;1710: 113– 121.

36. Coustou V, Besteiro S, Rivière L, Biran M, Biteau N, Franconi JM, et al. A mitochondrial NADH-dependent fumarate reductase involved in the production of succinate excreted by procyclic Trypanosoma brucei. J Biol Chem. 2005;280: 16559–16570.

37. Jitrapakdee S, St Maurice M, Rayment I, Cleland WW, Wallace JC, Attwood P V. Structure, mechanism and regulation of pyruvate carboxylase. Biochem J. 2008;413: 369–387.

38. Leger MM, Eme L, Hug LA, Roger AJ. Novel hydrogenosomes in the microaerophilic jakobid Stygiella incarcerata. Mol Biol Evol. 2016/06/08. 2016;33: 2318–2336.

39. Kikuchi G. The glycine cleavage system: composition, reaction mechanism, and physiological significance. Mol Cell Biochem. 1973;1: 169–187.

40. Dale RA. Catabolism of threonine in mammals by coupling of L-threonine 3-dehydrogenase with 2-amino-3-oxobutyrate-CoA ligase. Biochim Biophys Acta. 1978;544: 496–503.

41. Schertl P, Braun H-P. Respiratory electron transfer pathways in plant mitochondria. Front Plant Sci. 2014;5: 163.

42. Thomson JM, Gaucher EA, Burgan MF, De Kee DW, Li T, Aris JP, et al. Resurrecting ancestral alcohol dehydrogenases from yeast. Nat Genet. 2005;37: 630–635.

43. Hügler M, Wirsen CO, Fuchs G, Taylor CD, Sievert SM. Evidence for autotrophic CO_2_ fixation via the reductive tricarboxylic acid cycle by members of the ε subdivision of proteobacteria. J Bacteriol. 2005;187: 3020–3027.

44. Jiang L, Boufersaoui A, Yang C, Ko B, Rakheja D, Guevara G, et al. Quantitative metabolic flux analysis reveals an unconventional pathway of fatty acid synthesis in cancer cells deficient for the mitochondrial citrate transport protein. Metab Eng. 2017;43: 198–207.

45. Verschueren KHG, Blanchet C, Felix J, Dansercoer A, De Vos D, Bloch Y, et al. Structure of ATP citrate lyase and the origin of citrate synthase in the Krebs cycle. Nature. 2019;568: 571–575.

46. Taylor EB. Functional properties of the mitochondrial carrier system. Trends Cell Biol. 2017;27: 633–644.

47. Jackson JB, Peake SJ, White SA. Structure and mechanism of proton-translocating transhydrogenase. FEBS Lett. 1999;464: 1–8.

48. Yu Y, Samuelson J. Primary structure of an Entamoeba histolytica nicotinamide nucleotide transhydrogenase. Mol Biochem Parasitol. 1994;68: 323–328.

49. Yousuf MA, Mi-ichi F, Nakada-Tsukui K, Nozaki T. Localization and targeting of an unusual pyridine nucleotide transhydrogenase in Entamoeba histolytica. Eukaryot Cell. 2010/04/09. 2010;9: 926–933.

50. Kuchenreuther JM, Britt RD, Swartz JR. New insights into [FeFe] hydrogenase activation and maturase function. PLoS One. 2012;7: e45850.

51. Tachezy J, Doležal P. Iron–sulfur proteins and iron–sulfur cluster assembly in organisms with hydrogenosomes and mitosomes. In: Martin WF, Müller M, editors. Origin of Mitochondria and Hydrogenosomes. Berlin, Heidelberg: Springer Berlin Heidelberg; 2007. pp. 105–133.

52. Stairs CW, Eme L, Brown MW, Mutsaers C, Susko E, Dellaire G, et al. A SUF Fe-S cluster biogenesis system in the mitochondrion-related organelles of the anaerobic protist Pygsuia. Curr Biol. 2014;24: 1176–1186.

53. Thomé R, Gust A, Toci R, Mendel R, Bittner F, Magalon A, et al. A sulfurtransferase is essential for activity of formate dehydrogenases in Escherichia coli. J Biol Chem. 2012;287: 4671–4678.

54. Mansy SS, Wu G, Surerus KK, Cowan JA. Iron-sulfur cluster biosynthesis. Thermatoga maritima IscU is a structured iron-sulfur cluster assembly protein. J Biol Chem. 2002;277: 21397–21404.

55. Kaiser JT, Clausen T, Bourenkow GP, Bartunik HD, Steinbacher S, Huber R. Crystal structure of a NifS-like protein from Thermotoga maritima: implications for iron sulphur cluster assembly. J Mol Biol. 2000;297: 451–464.

56. Mi-ichi F, Makiuchi T, Furukawa A, Sato D, Nozaki T. Sulfate activation in mitosomes plays an important role in the proliferation of Entamoeba histolytica. PLoS Negl Trop Dis. 2011;5: e1263.

57. Mi-ichi F, Nozawa A, Yoshida H, Tozawa Y, Nozaki T. Evidence that the Entamoeba histolytica mitochondrial carrier family links mitosomal and cytosolic pathways through exchange of 3IZ-phosphoadenosine 5IZ-phosphosulfate and ATP. Eukaryot Cell. 2015;14: 1144–1150.

58. Liu S, He L, Yao K. The antioxidative function of alpha-ketoglutarate and its applications. Biomed Res Int. 2018;2018: 3408467.

59. Sonneborn TM. Methods in the general biology and genetics of Paramecium aurelia. J Exp Zool. 1950;113: 87–147.

60. Trager W. The cultivation of a cellulose-digesting flagellate, Trichomonas termopsidis, and of certain other termite protozoa. Biol Bull. 1934;66: 182–190.

61. Picelli S, Faridani OR, Björklund AK, Winberg G, Sagasser S, Sandberg R. Full-length RNA-seq from single cells using Smart-seq2. Nat Protoc. 2014;9: 171–181.

62. Bankevich A, Nurk S, Antipov D, Gurevich AA, Dvorkin M, Kulikov AS, et al. SPAdes: a new genome assembly algorithm and its applications to single-cell sequencing. J Comput Biol. 2012/04/16. 2012;19: 455–477.

63. Dick GJ, Andersson AF, Baker BJ, Simmons SL, Thomas BC, Yelton AP, et al. Community-wide analysis of microbial genome sequence signatures. Genome Biol. 2009/08/21. 2009;10: R85–R85.

64. Treitli SC, Kolisko M, Husník F, Keeling PJ, Hampl V. Revealing the metabolic capacity of Streblomastix strix and its bacterial symbionts using single-cell metagenomics. Proc Natl Acad Sci U S A. 2019;116: 19675–19684.

65. Zhu B-H, Xiao J, Xue W, Xu G-C, Sun M-Y, Li J-T. P_RNA_scaffolder: a fast and accurate genome scaffolder using paired-end RNA-sequencing reads. BMC Genomics. 2018;19: 175.

66. Stanke M, Schöffmann O, Morgenstern B, Waack S. Gene prediction in eukaryotes with a generalized hidden Markov model that uses hints from external sources. BMC Bioinformatics. 2006;7: 62.

67. Haas BJ, Salzberg SL, Zhu W, Pertea M, Allen JE, Orvis J, et al. Automated eukaryotic gene structure annotation using EVidenceModeler and the Program to Assemble Spliced Alignments. Genome Biol. 2008;9: R7–R7.

68. Grabherr MG, Haas BJ, Yassour M, Levin JZ, Thompson DA, Amit I, et al. Full-length transcriptome assembly from RNA-Seq data without a reference genome. Nat Biotechnol. 2011/05/17. 2011;29: 644–652.

69. Langmead B, Salzberg SL. Fast gapped-read alignment with Bowtie 2. Nat Methods. 2012/03/06. 2012;9: 357–359.

70. Kim D, Paggi JM, Park C, Bennett C, Salzberg SL. Graph-based genome alignment and genotyping with HISAT2 and HISAT-genotype. Nat Biotechnol. 2019;37: 907–915.

71. Simão FA, Waterhouse RM, Ioannidis P, Kriventseva E V, Zdobnov EM. BUSCO: assessing genome assembly and annotation completeness with single-copy orthologs. Bioinformatics. 2015/06/11. 2015;31: 3210–3212.

72. Altschul SF, Gish W, Miller W, Myers EW, Lipman DJ. Basic local alignment search tool. J Mol Biol. 1990;215: 403–410.

73. Eddy SR. A new generation of homology search tools based on probabilistic inference. Genome Inf. 2010/02/25. 2009;23: 205–211.

74. Jones P, Binns D, Chang HY, Fraser M, Li W, McAnulla C, et al. InterProScan 5: genome-scale protein function classification. Bioinformatics. 2014/01/24. 2014;30: 1236–1240.

75. Kearse M, Moir R, Wilson A, Stones-Havas S, Cheung M, Sturrock S, et al. Geneious Basic: an integrated and extendable desktop software platform for the organization and analysis of sequence data. Bioinformatics. 2012/05/01. 2012;28: 1647–1649.

76. Almagro Armenteros JJ, Salvatore M, Emanuelsson O, Winther O, von Heijne G, Elofsson A, et al. Detecting sequence signals in targeting peptides using deep learning. Life Sci Alliance. 2019;2: e201900429.

77. Horton P, Nakai K. Better prediction of protein cellular localization sites with the k nearest neighbors classifier. Proc Int Conf Intell Syst Mol Biol. 1997;5: 147–152.

78. Blum T, Briesemeister S, Kohlbacher O. MultiLoc2: Integrating phylogeny and gene ontology terms improves subcellular protein localization prediction. BMC Bioinformatics. 2009/09/03. 2009;10: 274.

79. Kume K, Amagasa T, Hashimoto T, Kitagawa H. NommPred: Prediction of mitochondrial and mitochondrion-related organelle proteins of nonmodel organisms. Evol Bioinform Online. 2018;14: 1176934318819835.

80. Reumann S. Specification of the peroxisome targeting signals type 1 and type 2 of plant peroxisomes by bioinformatics analyses. Plant Physiol. 2004;135: 783–800.

81. Krogh A, Larsson B, von Heijne G, Sonnhammer EL. Predicting transmembrane protein topology with a hidden Markov model: application to complete genomes. J Mol Biol. 2001/01/12. 2001;305: 567–580.

82. Neuberger G, Maurer-Stroh S, Eisenhaber B, Hartig A, Eisenhaber F. Prediction of peroxisomal targeting signal 1 containing proteins from amino acid sequence. J Mol Biol. 2003;328: 581–592.

83. Katoh K, Standley DM. MAFFT multiple sequence alignment software version 7: improvements in performance and usability. Mol Biol Evol. 2013/01/19. 2013;30: 772– 780.

84. Hall TA. BioEdit: a user-friendly biological sequence alignment editor and analysis program for Windows 95/98/ NT. Nucleic Acids Symp Ser. 1999;41: 95–98.

85. Stamatakis A. RAxML version 8: A tool for phylogenetic analysis and post-analysis of large phylogenies. Bioinformatics. 2014;30: 1312–1313.

86. Capella-Gutiérrez S, Silla-Martínez JM, Gabaldón T. trimAl: a tool for automated alignment trimming in large-scale phylogenetic analyses. Bioinformatics. 2009/06/10. 2009;25: 1972–1973.

87. Nguyen LT, Schmidt HA, von Haeseler A, Minh BQ. IQ-TREE: a fast and effective stochastic algorithm for estimating maximum-likelihood phylogenies. Mol Biol Evol. 2014/11/06. 2015;32: 268–274.

88. Wang H-C, Minh BQ, Susko E, Roger AJ. Modeling site heterogeneity with posterior mean site frequency profiles accelerates accurate phylogenomic estimation. Syst Biol. 2018;67: 216–235.

89. Hoang DT, Chernomor O, von Haeseler A, Minh BQ, Vinh LS. UFBoot2: Improving the ultrafast bootstrap approximation. Mol Biol Evol. 2017;35: 518–522.

90. Gietz RD, Woods RA. Yeast transformation by the LiAc/SS Carrier DNA/PEG method. In: Xiao W, editor. Yeast Protocols. Totowa, NJ: Humana Press; 2006. pp. 107–120.

