## Supplementary Figures for "Anaerobic derivates of mitochondria and peroxisomes in the free-living amoeba *Pelomyxa schiedti* revealed by single-cell genomics"

**S1 – S6 FIGURES**

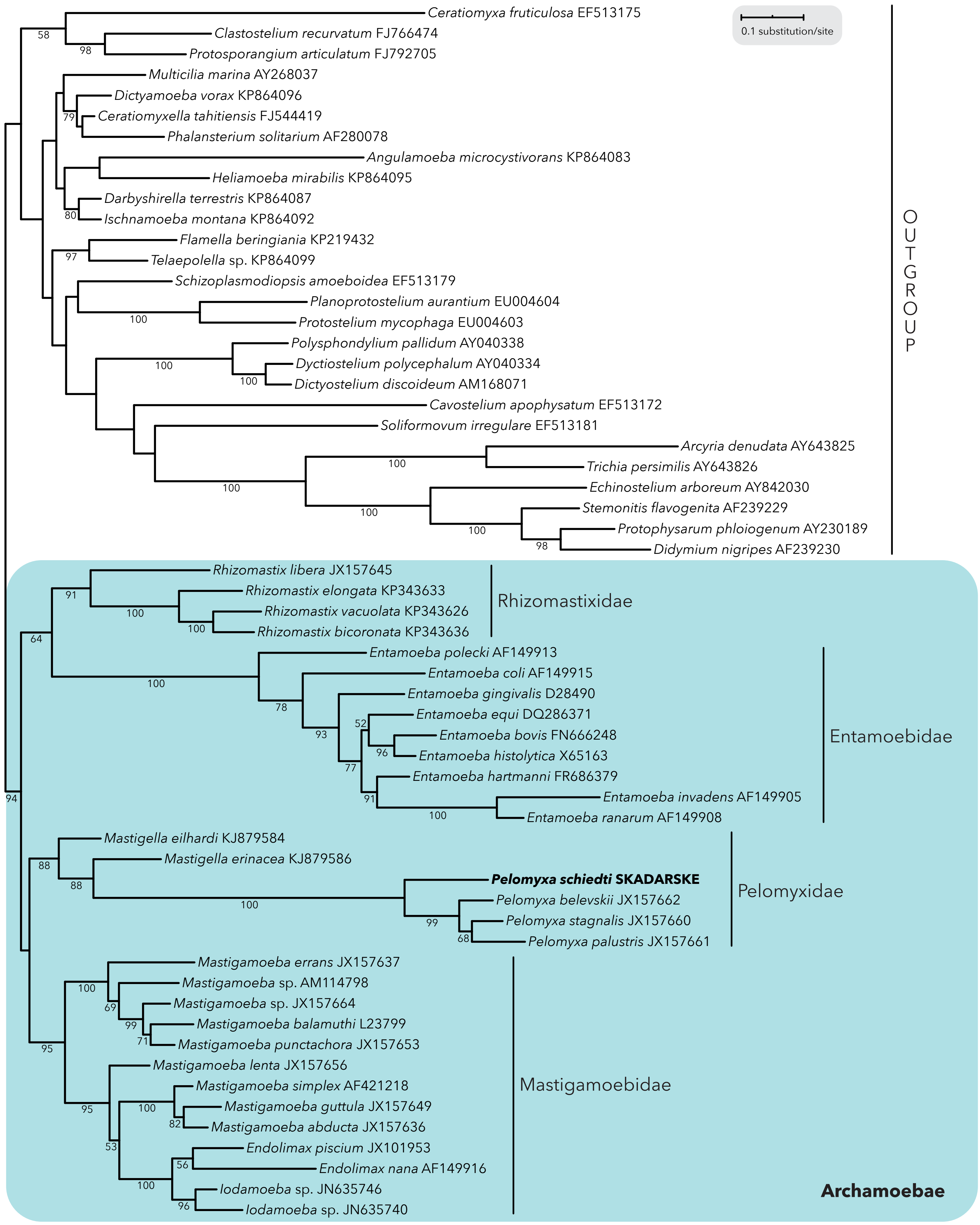


**S1 Fig. Phylogenetic analysis of amoebozoan 18S rDNA.** The Maximum Likelihood tree places *Pelomyxa schiedti* in monophyletic Pelomyxidae group inside monophyletic Archamoebae. Standard bootstrap support values are shown when ≥50%.

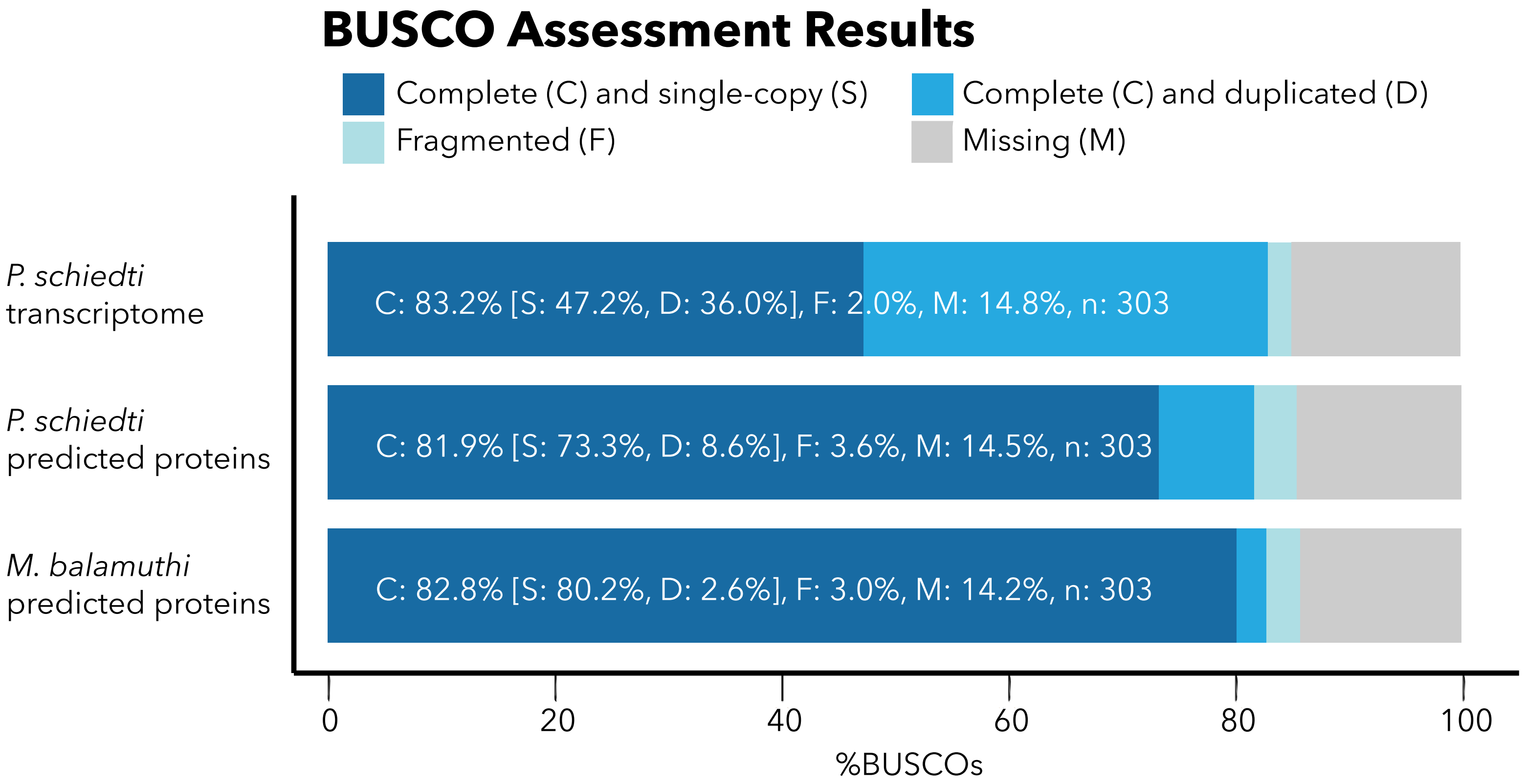


**S2 Fig. BUSCO analysis of the *Pelomyxa schiedti* transcriptome and predicted proteins.** Completeness of *P. schiedti* datasets were assessed using the odb9_eukaryota dataset and compared with completeness of predicted proteins from *Mastigamoeba balamuthi*.

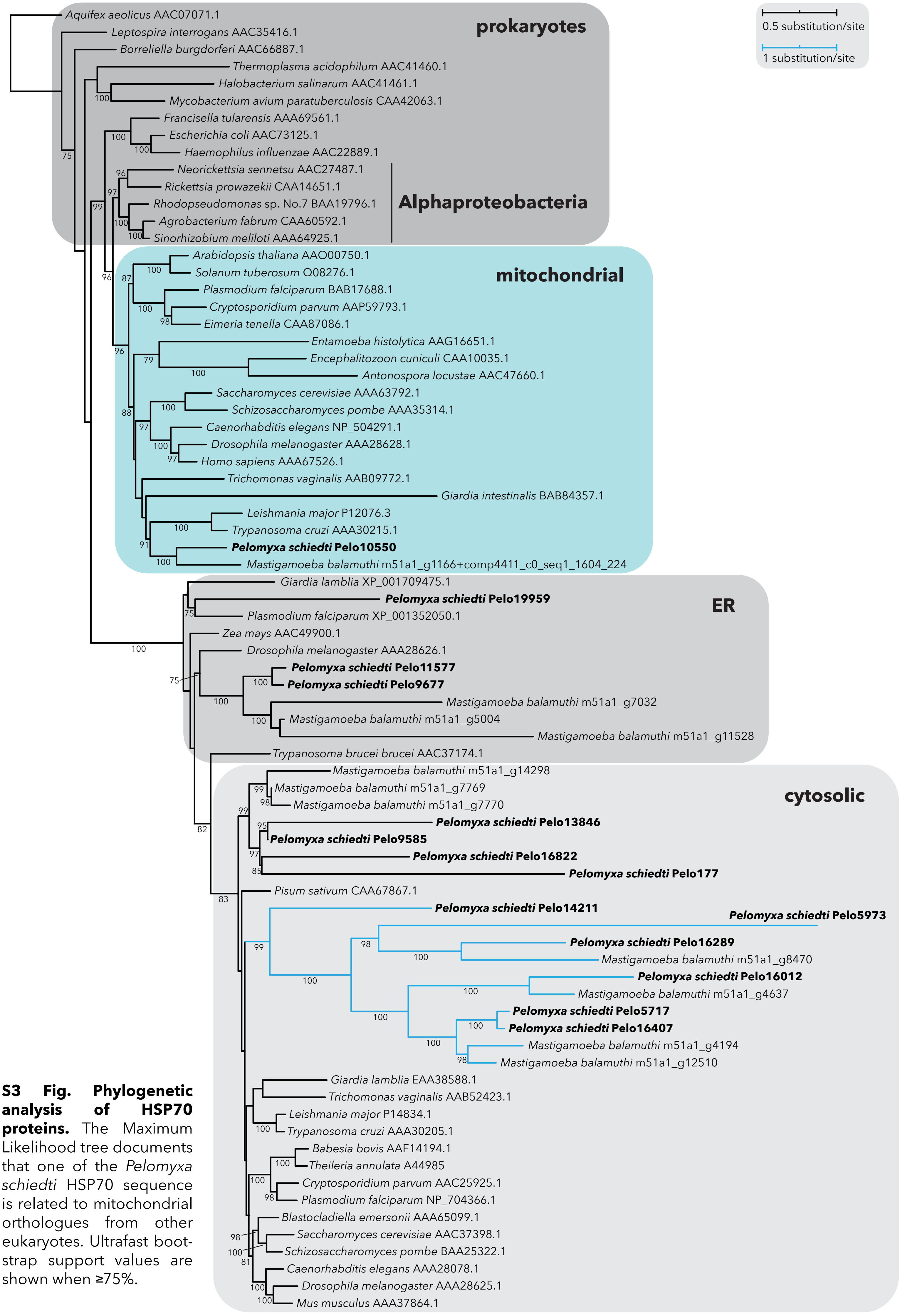


**S3 Fig. Phylogenetic analysis of HSP70 proteins.** The Maximum Likelihood tree documents that one of the *Pelomyxa schiedti* HSP70 sequence is related to mitochondrial orthologues from other eukaryotes. Ultrafast bootstrap support values are shown when ≥75%.

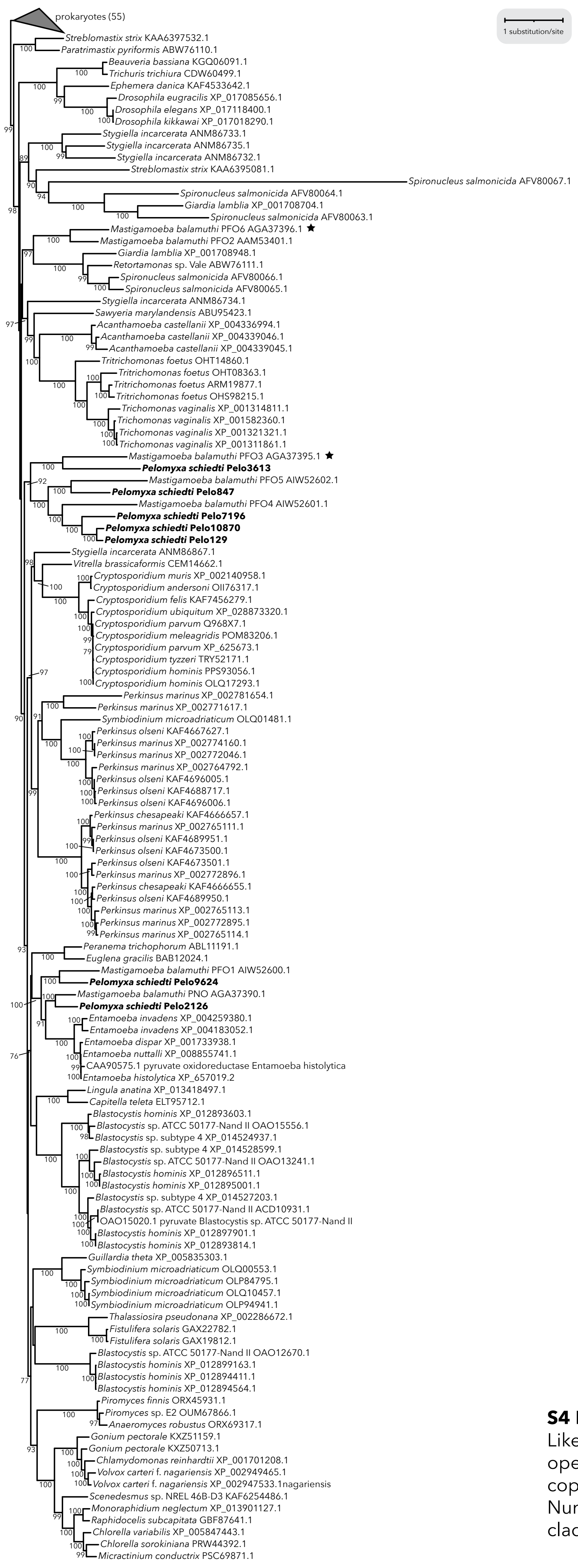


**S4 Fig. Phylogenetic analysis of PFO enzymes.** The Maximum Likelihood phylogenetic tree identified a PFO version putatively operating in *Pelomyxa schiedti* MRO. Hydrogenosomal PFO copies of *Mastigamoeba balamuthi* are marked with stars. Number in parenthesis shows number of species in the collapsed clade. Ultrafast bootstrap support values are shown when ≥75%.

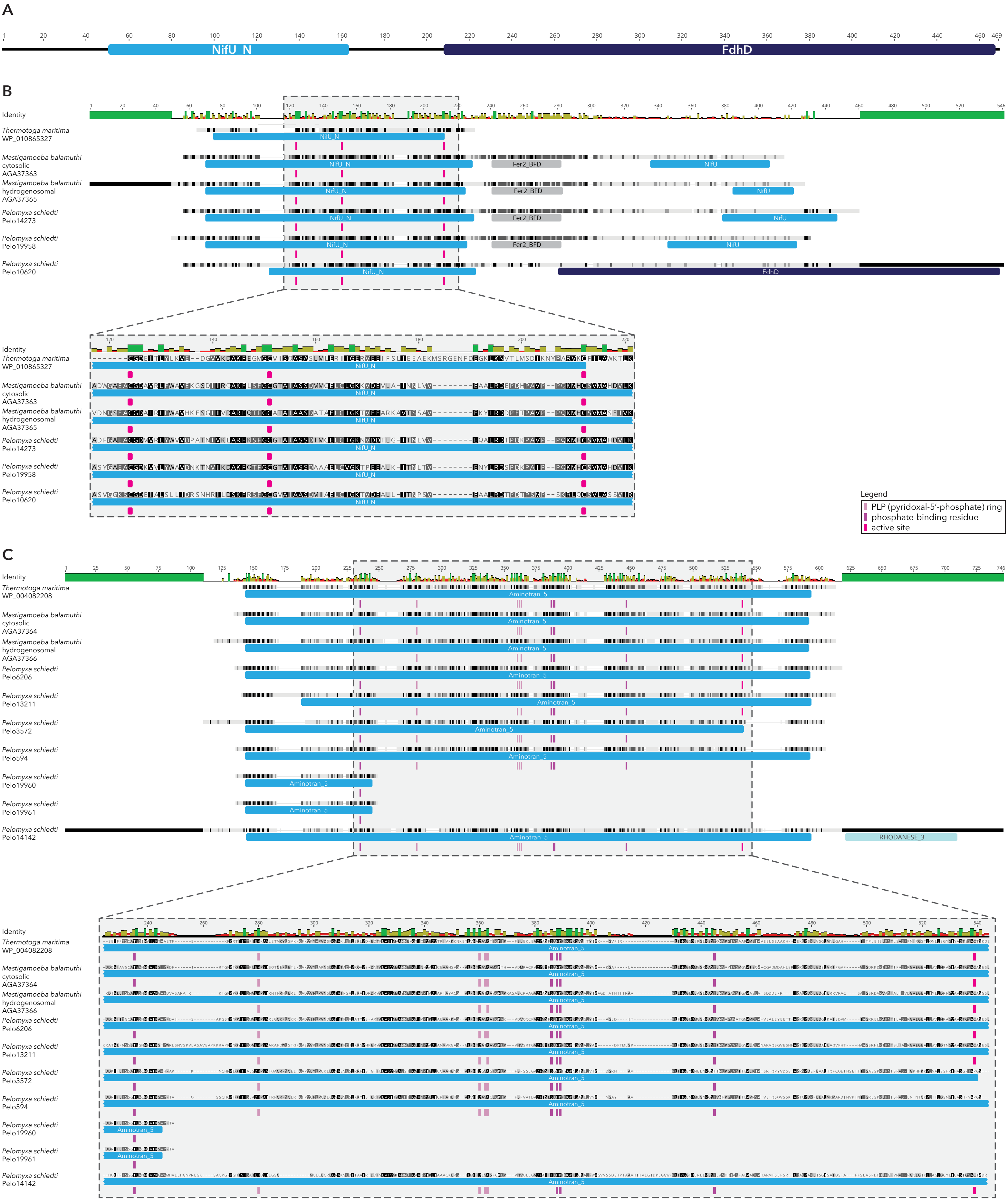


**S5 Fig. Sequences of *Pelomyxa schiedti* components of NIF system.** (A) The diagram depicts *P. schiedti* protein Pelo10620 composed of a NifU N-terminal domain fused to a FdhD (formate dehydrogenase accessory sulfurtransferase) C-terminal domain as determined by InterProScan. (B-C) Sequence alignment of NifU (B) and NifS (C) proteins from *P. schiedti* and *Mastigamoeba balamuthi* in comparison with bacterial homologues from *Thermotoga maritima*. The amino acid residues necessary for the function of NifU and NifS are labeled according to the legend.

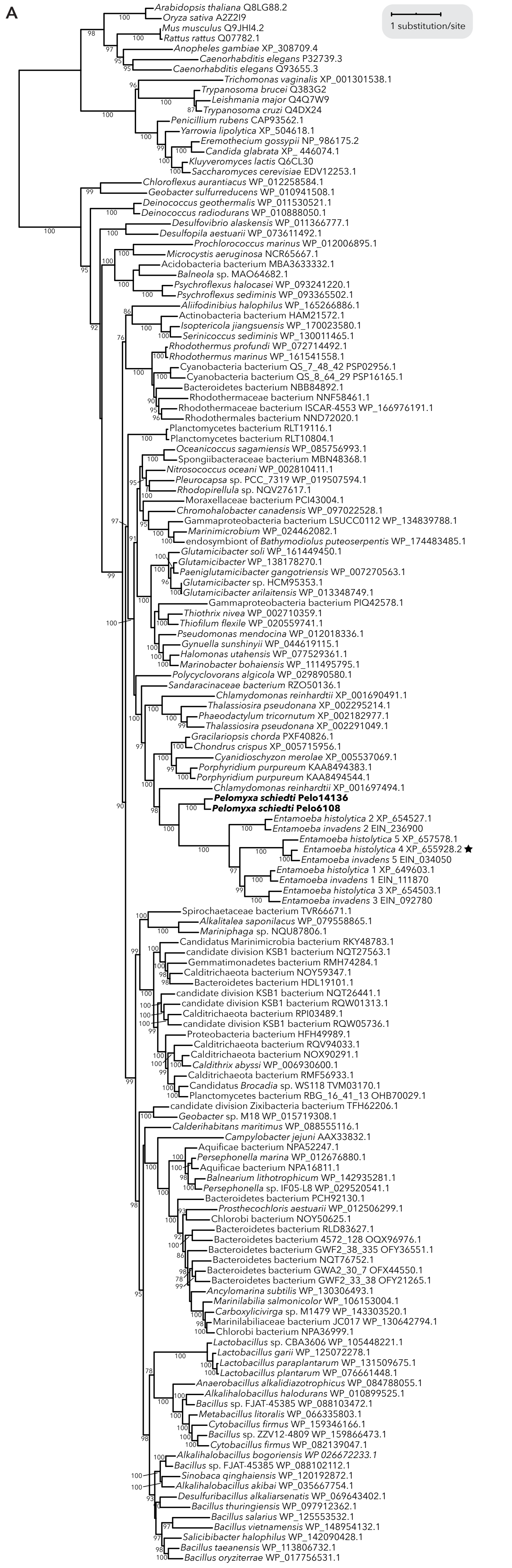


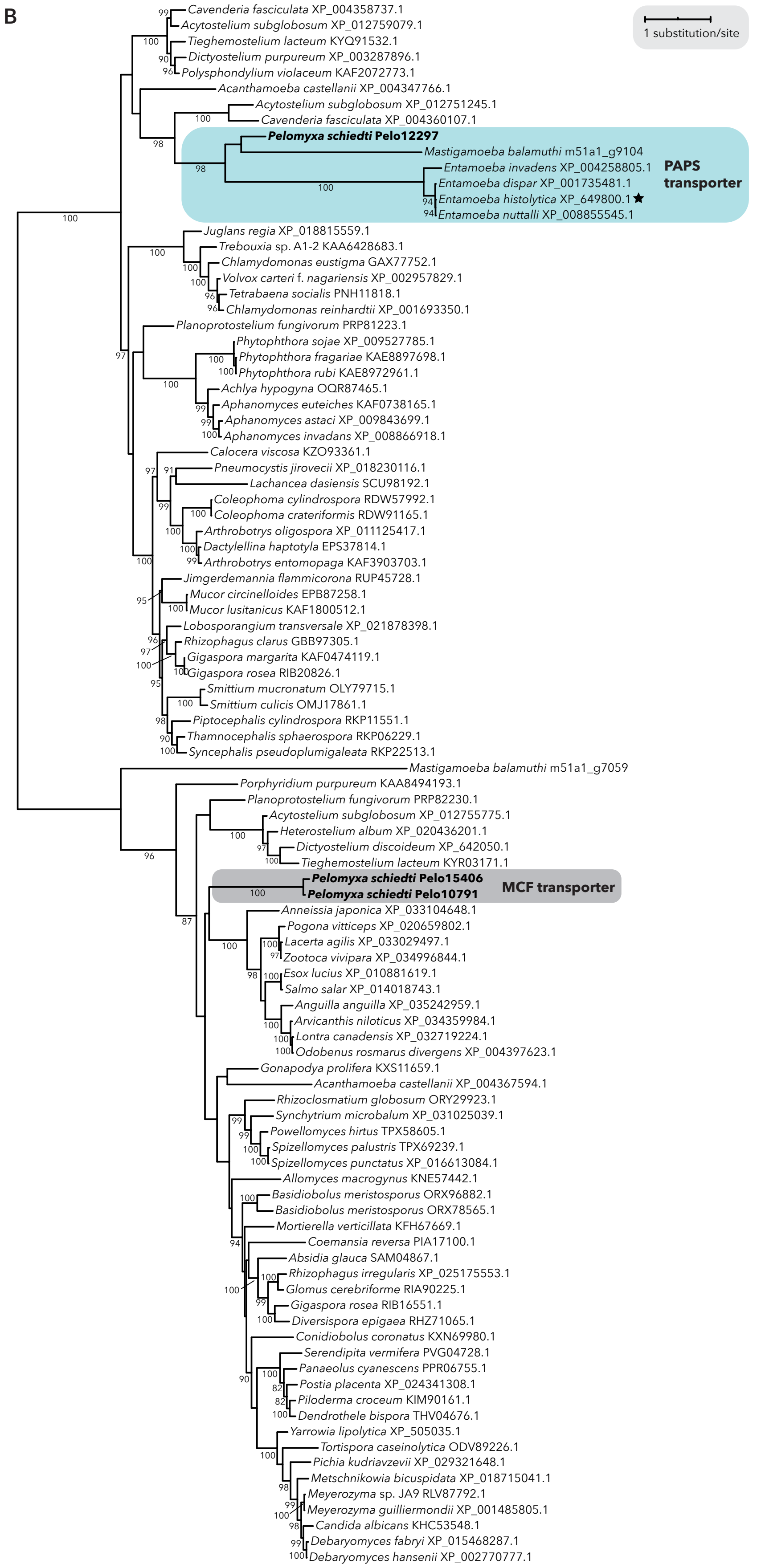


**S6 Fig. Phylogenetic analysis of transporters involved in the sulfate activation pathway.** (A) The phylogenetic analysis did not resolve which one of the sodium/sulfate symporters of *Pelomyxa schiedti* is related to the *Entamoeba histolytica* mitosomal transporter. (B) The Maximum Likelihood phylogenetic tree confirms one *P. schiedti* transporter as PAPS (3′-phosphoadenosine 5′-phosphosulfate) transporter, while two others belong to a broader mitochondrial carrier family of transporters. Experimentally proven mitosomal transporters of *E. histolytica* are marked with stars. Ultrafast bootstrap support values are shown when ≥75%.
